# Integration of an invasive plant in hummingbird and flower mite networks is driven by ecological fitting and generalization

**DOI:** 10.1101/2024.12.20.629806

**Authors:** Carlos García-Robledo, Joel Alvarado

## Abstract

Most plant communities worldwide include exotic plants, which did not evolve with local organisms. The central goal of this study is to test if native organisms expanding to novel hosts are usually generalists or specialists. Here we studied new association between hummingbirds, flower mites and *Musa velutina* (Musaceae), an exotic plant native to Asia currently invading lowland forests in Costa Rica. Hummingbirds are pollinators, but flower mites feed on nectar without contributing to pollen transfer. Flower mites hitch rides on hummingbird beaks to colonize new flowers. To determine the original diet breadth of hummingbird and flower mite species, we assembled interaction networks including 33 native host plants, 14 hummingbird and 19 flower mite species. Transportation analyses show that flower mites’ colonization of native or exotic hosts is not constrained by hummingbird flight connections. We identified four hummingbird species visiting *Musa velutina.* DNA barcodes analyses identified only one species of flower mite colonizing flowers of *M. velutina*. All new associations with *M. velutina* involved generalist hummingbird and flower mite species. *Musa velutina* displays both male and female flowers. Although flowers of both sexes were equally visited by hummingbirds, mites were 15 times more abundant in male than in female flowers. This might be the result of constant immigration coupled with population growth. Only half of the mites hitching rides on hummingbird beaks emigrate to newly open flowers. Our results show that *M. velutina* integration to a plant community occurs mainly by establishing interactions with generalists.

---

The introduction of exotic species is affecting the structure and composition of native ecosystems worldwide. Exotic plants may also affect functional ecosystem attributes, for example, the structure of plant - pollinator networks (Frost et al., 2019). Traits evolved by exotic plants in their original habitats may facilitate the assembly of novel interactions in the novel habitat. This complementarity of not coevolved traits is known as ecological fitting (Agosta & Klemens, 2008; Janzen, 1985). Novel interactions between exotic plants and native organisms, besides affecting the life history of native organisms, also represent an opportunity to understand processes involved in the integration of novel species in plant communities.

One prediction of ecological fitting is that native generalists may have an advantage over specialists to exploit novel host plants (Agosta & Klemens, 2008). Most studies report generalist pollinators interacting with exotic plants (Lopezaraiza–Mikel, Hayes, Whalley, & Memmott, 2007; Memmott & Waser, 2002). However, empirical studies also support the opposite prediction. In insect pollinator networks, specialist insects are more likely to interact with exotic plants than generalists (Stouffer, Cirtwill, & Bascompte, 2014).

In the neotropics, hummingbirds (Family Trochilidae) are a major group of pollinators (Barreto et al., 2024). Many exotic plants introduced to the neotropics display convergent traits associated with hummingbird pollination such as tubular corollas and flowers of bright colors (Sánchez & Lara, 2024). It is common for hummingbirds to visit exotic plants as soon as they learn that these plants are a reliable source of nectar (Sánchez & Lara, 2024).

Hummingbirds are also associated with a group of neotropical floral parasites, the hummingbird flower mites (Acari: Mesostigmata: Ascidae) (Baker & Yunker, 1964). This group of mites are found feeding inside the flowers of hummingbird-pollinated plants (Colwell, 1973). Flower mites feed on nectar and pollen without apparently contributing to pollination. This guild of tiny nectar thieves depends on hummingbirds for transportation. Flower mites hitch rides on hummingbird beaks to eventually disembark in newly open flowers (Baker & Yunker, 1964; Colwell, 1973). Although hummingbird-mite interactions were usually described as a commensalism (Colwell, 1986), more recent studies show that flower mites may consume more than half of the nectar produced in a flower, potentially competing with hummingbirds (Colwell, 1995; Da Cruz, Righetti De Abreu, & Van Sluys, 2007; Lara & Ornelas, 2001; Paciorek, Moyer, Levin, & Halpern, 1995).

This study focuses on two guilds of hummingbirds and flower mites at La Selva Biological Station, a tropical lowland forest in Costa Rica. In our study site we observed hummingbirds visiting flowers of the pink banana (*Musa velutina* H. Wendl. & Drude, Musaceae, Zingiberales). This plant is originally from Asia, and after its introduction to Costa Rica, it became one of the most aggressive invasive plants in lowland and montane forests (Valverde, 2013).

The central goal of this study is to determine if the integration of *Musa velutina* into local pollination and flower parasite networks is facilitated by generalist or specialist species. We are particularly interested in the process of colonization of *M. velutina* by flower mites, as it might be limited by the patterns of flower visitation by hummingbirds. Using the analogy of hummingbird species as “flight connections” between plants, if *Musa velutina* is visited by generalist hummingbirds, these generalized flight connections will increase the connectivity with other plant species. A more diverse pool of flight connections may facilitate the colonization of *M. velutina* by more flower mite species. If *M. velutina* is visited by specialist hummingbirds, fewer flight connections may limit mite transportation to the novel host. An alternative is that flower mites are very specialized to their original hosts, and flower mites will not colonize *M. velutina*, *Musa velutina* is monoecious, producing at first female flowers, which only offer nectar. After a few days, plants shift to produce male flowers, which offer both nectar and pollen. We are interested in determining if flower mites prefer flowers only offering nectar, or flowers offering a mix of nectar and pollen.

To determine if generalist or specialist pollinators and nectar parasites are expanding their diet to *Musa velutina*, our first step was to determine pollinator and nectar parasite diet breadths. To determine host plant use for each species, we assembled the hummingbird and flower mite interaction networks at La Selva. Using network analyses, we determined if generalists or specialists are colonizing the novel host plant. To determine if flower mite transportation is limited by the available hummingbird flight routes, first we assessed how connected are native plants by estimating the number of shared hummingbird species. We also assessed how connected is each native plant species to *Musa velutina* by estimating the number of hummingbird species shared between the novel host and each native plant.

Finally, we estimated for each flower mite species the number of flight routes from their native hosts to the novel host plant. By combining these results, we determined if colonization of *M. velutina* by flower mites is limited by available flight routes, or by intrinsic preferences of generalist and specialist floral parasites encountering an exotic plant.

## METHODS

### Study site and species

We performed this study at La Selva Biological Station (10° 25′ 19.2″ N, 84° 0′ 54″ W), a tropical rain forest in the Caribbean slope of Costa Rica (McClearn et al., 2016). Elevation at La Selva ranges from 35 to 137 m.a.s.l. Rainfall varies from 152.0 mm in March to 480.7 mm in July (McDade, 1994). The average annual temperature is 23.6 °C (McDade, 1994).

At La Selva, the flowers of at least 32 plant species are visited by 14 species of hummingbirds (Supplement S1). Inside the flowers, hummingbird flower mites feed on pollen and nectar, and hitch rides on hummingbird beaks to colonize new flowers (Baker & Yunker, 1964; Colwell, 1973).

Previous studies incorrectly reported that hummingbird flower mites are extremely specialized, and a single mite species is usually associated with each host plant (Colwell, 1986). Using the DNA barcode CO1, we discovered that flower mites in many cases are generalists, and flowers usually host a mix of generalist and specialist flower mite species (Bizzarri, Baer, & García-Robledo, 2022, 2023; G. García Franco, Martinez Burgoa, & Pérez, 2001; Kress, 2022). At La Selva, 19 hummingbird flower mite species visit 18 plant species (Supplement S1).

*Musa velutina* H. Wendl. & Drude (Musaceae), also known as the “pink banana” is native to Assam and the eastern Himalayas (Govaerts, Nic Lughadha, Black, Turner, & Paton, 2021). *Musa velutina* was initially introduced to Trinidad in 1938 (Cheesman, 1949). This species was documented for the first time in Costa Rica in 1987, at a locality near La Selva Biological Station (Avalos, Chacón Madrigal, & Artavia Rodríguez, 2021). *Musa velutina* is considered an aggressive invasive species in tropical lowland and montane forests in Costa Rica (Zuchowski & Forsyth, 2007). At La Selva, *M. velutina* is currently invading primary and secondary forests, usually in areas close to rivers (Balderama, Schoenberg, Murray, & Rundel, 2011).

*Musa velutina* produces a single erect inflorescence. Bracts subtend groups of flowers, known as banana “hands”. Each morning, a pink bract opens to expose a set of flowers. This species of wild banana is monoecious, with female and male flowers produced in the same inflorescence (Kirchoff, 2017). At La Selva, we observed that plants first produce 1 to 6 female flower hands, and transition to produce 7 to 40 male hands. Female and male hands have a similar number of flowers (Mean = 6 flowers, Min = 3, Max = 13, N = 28). Inflorescences produce flowers for about a month (Min = 11 d, Max = 49 d Mean = 29.1 d, N = 21).

In its native range, *Musa velutina* is pollinated by bats (Percival, 1979). Female and male flowers produce nectar with similar concentration, *ca*. 15%. However, female flowers produce less nectar than male flowers (Percival, 1979). In Costa Rica, inflorescences are visited by hummingbirds and stingless bees in the genus *Trigona* (Valverde, 2013).

### Hummingbird interactions with native plants and *Musa velutina*

Hummingbird-plant interactions had been studied at La Selva for the last fifty years (Supplement S1). To determine which hummingbird species visit each native plant species in our study site, we assembled a hummingbird-plant interactions network by combining our observations with published interaction records (Supplement S1).

To determine which hummingbird species visit *Musa velutina*, we used high-resolution cameras (Canon PowerShot SX530 HS, Canon Inc. Tokyo, Japan). We modified the cameras for motion detection by installing in the SD cards the firmware Canon Hack Development Kit, available at http://chdk.wikia.com/wiki/CHDK (Juárez, Vargas, & Kay, 2023; Maguiña Conde, Zuñiga Rivas, & Kay, 2023; Steen, 2017). The cameras were placed at 3 to 4 m from inflorescences and recorded each visit for 5 seconds. We recorded visits from 6:00 AM to 1:00 PM.

### Interactions of hummingbird flower mites with native plants and *Musa velutina*

To determine the association between flower mite species and hummingbird-pollinated plants at La Selva, we collected flower mites from each native plant species. Mites were fixed in ETOH 95%. Individual mites were placed in 96 well plates. To identify mite species, we amplified the DNA barcode CO1 (Hebert, Cywinska, Ball, & DeWaard, 2003). We assembled the plant-flower mite network by combining new interaction records in this study with interactions reported in one of our previous studies (see methods in Bizzarri et al., 2022, 2023). All sequences were deposited in GenBank (Accession Numbers. MW14554-MW147005 and PQ438945-PQ439121).

To determine which mite species are expanding their diets to *Musa velutina,* we collected flower mites from recently open flowers (see sample size in the results section). Mites were fixed in ETOH 95%, then sequenced to obtain the DNA barcode CO1. We identified the mite species colonizing *Musa velutina* by comparing mite CO1 sequences of individuals collected in *M. velutina* with our DNA barcode library. Sequence comparison was performed using the BLAST algorithm. (Camacho et al., 2009). All analyses were performed using the program Geneious (Geneious-Prime-2023.2.1, 2023).

### Changes in network structure after diet expansions to *Musa velutina*

Hummingbird and flower mite associations with their host plants usually display nested network structures (Vizentin-Bugoni et al., 2016). This structure emerges from some plants being visited by mixes of generalist and specialist species, but plants visited by a single species usually interact with generalists. An increase in nestedness is associated with higher resilience to species coextinction (Duan, Zhai, Hou, Zhou, & Rong, 2023).

To determine changes in nestedness in hummingbird and flower mite communities at La Selva, we first calculated nestedness in networks including only native plants (hereafter original networks). We also calculated nestedness for networks including *M. velutina* (hereafter novel networks). Absolute nestedness values are expected to decrease in the novel networks if the species expanding their diets are specialists (Duan et al., 2023). If generalist species are expanding their diets to *M. velutina*, we expect an increase in nestedness in novel networks (Duan et al., 2023).

For hummingbird and mite networks, we estimated nestedness using the metric NODF (Nestedness Overlap and Decreassing Fill, Almeida Neto, Guimaraes, Guimaraes Jr, Loyola, & Ulrich, 2008). NODF values range from 0 (not nested) to 100 (all interactions fit below the matrix diagonal, *i.e.*, perfectly nested). The NODF metric corrects for matrix fill and matrix dimensions, making possible to compare nestedness between matrices with different dimensions (Almeida Neto et al., 2008).

To determine differences in nestedness when including hummingbird or mite interactions with *M. velutina,* we performed permutation tests between original and novel matrices. For all matrices, we performed 1000 permutations. We estimated NODF for all matrices using the function NODF2 in the package bipartite in program R (Almeida Neto et al., 2008; Dormann, Fründ, Blüthgen, & Gruber, 2009). Probability values were calculated based on the histogram of native vs novel matrix NODF differences. All analyses were performed using the package bipartite in Program R and Rstudio IDE (Dormann et al., 2009; R_Core_Team, 2024; RStudio_Team, 2024).

### Flight connections between native plants and *Musa velutina*

To determine how accessible is *Musa velutina* to each mite species from their native hosts, we calculated the number of hummingbird species connecting each plant species pair (Figure 2). We estimated D*_i_*, the degree of connectivity for each plant species:

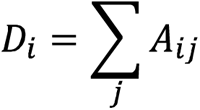

Where A is the quantitative matrix including the number of hummingbird species shared by plant species *i* and *j*. We sorted the matrix elements from higher to lower degree of connectivity. These analyses allowed us to determine if *M. velutina* is grouped with highly connected native plants, suggesting that many flower mite species have the potential to expand their diets to the novel host. An alternative is that *M. velutina* is visited by a few specialized hummingbirds, limiting the colonization by most mite species.

To determine the potential routes available for each mite species *i* from their host plants to *M. velutina*, we calculated for each element the plant-mite interaction matrix excluding *M. velutina* (Figure 1B), the number of hummingbird species *H* flying from native host *j* to *M. velutina*.

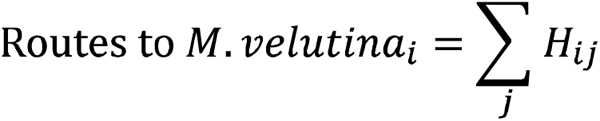

**Figure 1.**
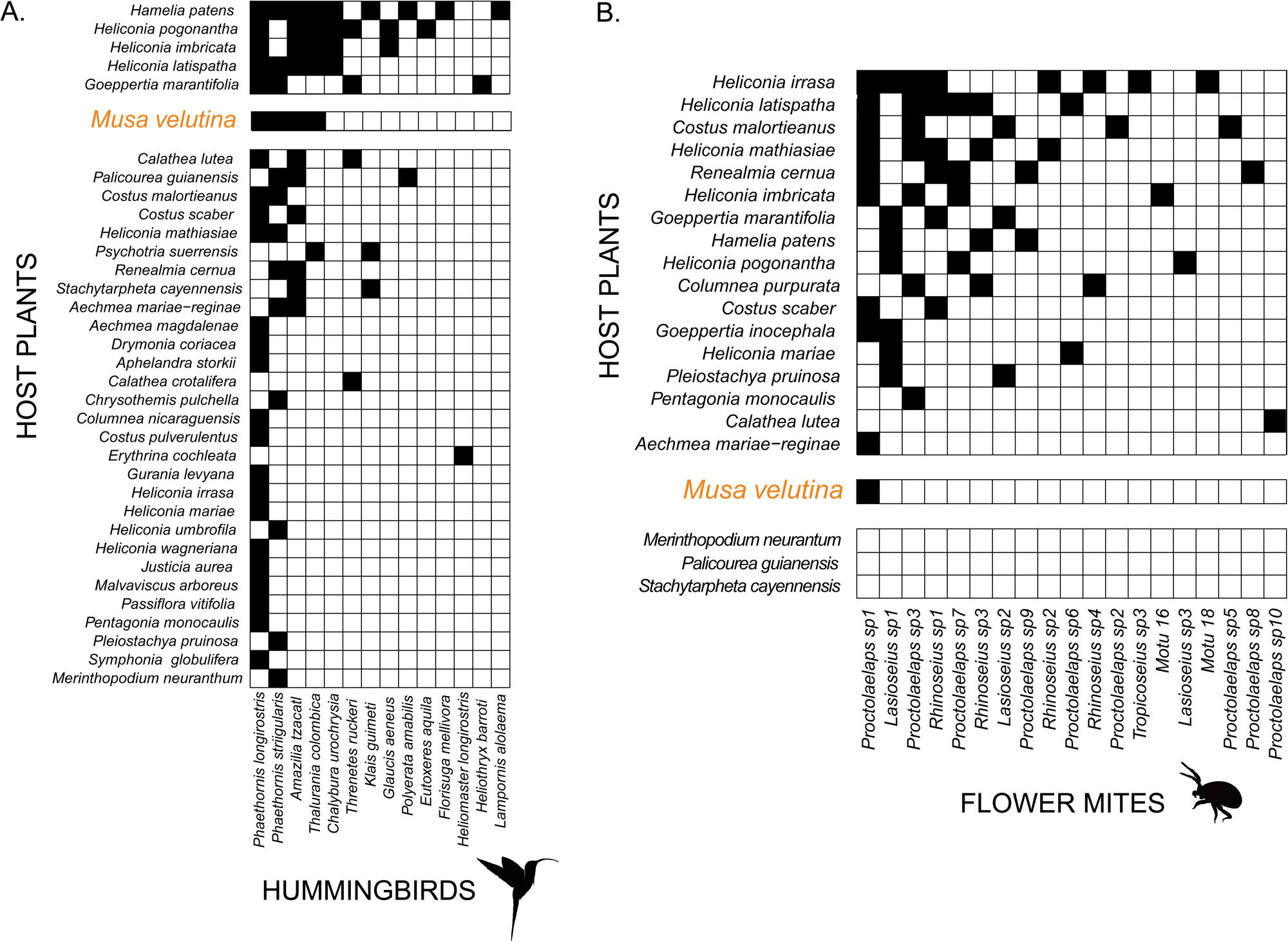
Interactions between plants, hummingbird and flower mites at La Selva Biological Station, Costa Rica. A. Plant-hummingbird network. Black cells represent hummingbird species visiting each host plant. Hummingbird species visiting *Musa velutina* are highlighted in orange. B. Plant-flower mite network. Note three plant species at the bottom, with no record of flower mites. The only mite species expanding its diet to *Musa velutina* is highlighted in orange. References for each plant-hummingbird interaction are included in Supplement S1.

The resulting frequency distribution represents the available flight routes for each mite species to colonize *M. velutina*. Using the flower mites collected in *M. velutina* identified using DNA barcoding, we determined if mite species with more flight connections are more likely to expand their diets to the novel host.

### Hummingbird visits to male and female *M. velutina* flowers

To determine differences in the number of hummingbird visits to female and male flower hands, we placed cameras at 3-4 m from the flowers. Cameras were set to record visits for five seconds. We recorded hummingbird visits from 6:00 AM to 1:00 PM. We only surveyed each inflorescence during either its female or male phases. To determine differences in the number of visits per flower hand, we performed a generalized linear model, including plant individuals and sex as factors, and number of hummingbird visits per hand as response variable.

### Hummingbird mite transit and colonization of male and female *M. velutina* flowers

To determine differences in mite colonization and migration from female and male *M. velutina* flowers, we counted the number of mites embarking or disembarking in each hummingbird visit. Videos were analyzed using the application Adobe Premiere Pro V24.1 (Adobe Inc. San Jose, CA, USA). Differences in the number of mites embarking or disembarking between male and female flowers were determined using a generalized lineal model, including hummingbird species, flower sex and transit (*ie*., embarking vs disembarking) as factors.

To determine differences in the number of mites present in female and male flowers, we collected individual flowers in ETOH 95%, then counted the number of flower mites per flower. We tested for differences in the average number of flower mites using a Welch Two Sample t-test (Welch, 1947).

## RESULTS

### Hummingbird interactions with native plants and *Musa velutina*

At La Selva, 14 hummingbird species visit 33 native plant species. The most generalist hummingbird species is *Phaethornis longirostris*. This hummingbird species visits 72% of the native species at La Selva. The second most generalist hummingbird species is *Phaethornis striigularis*. This species visits 31% of all native plant species included in this study. We observed *P. striigularis* feeding at the base of the corollas of *Costus malortieanus* and *Aechmea mariae-reginae*, therefore, it remains unknown if flower mites can use this hummingbird species for transportation to these host plants. *Phaethornis striigularis* also visits flowers of the bat pollinated plant *Merinthopodium neuranthum.* All other hummingbird species were recorded visiting between one and four host plants (Figure 1A).

After 200 hours of video, we observed the four most generalist hummingbird species visiting *M. velutina* (Figure 1A). We recorded 165 visits by *P. longirostris,* 137 visits by *A. tzacatl,* 24 visits by *P. striigularis* and two visits by *T. colombica.* All visits by *P. striigularis* were legitimate, suggesting that this hummingbird species contributes to flower mite transportation to the novel host (Figure 1A).

### Interactions of hummingbird flower mites with native plants and *Musa velutina*

Based on 1884 DNA barcode sequences, we identified 19 flower mite species interacting with 17 host plants (Figure 1B). The most generalist flower mite species is *Proctolaelaps* sp1 (Figure 1B). The diet breadth of all other flower mites ranges from generalist species using seven host plants, to specialist mites recorded in a single plant species (Figure 1B). Most plant species host a mix of generalist and specialist mite species (Figure 1B).

We sequenced 156 DNA barcodes from flower mites collected in 14 individuals of *M. velutina.* All individuals were identified as *Proctolaelaps* sp1, the most generalist flower mite species at La Selva (Figure 1B).

### Changes in network structure after diet expansions to *Musa velutina*

The NODF value for the hummingbird-plant network increased after including novel interactions with *M. velutina* (NODF_original_= 38.0, NODF_novel_ = 40.4). This result is not surprising, as *M. velutina* is visited by the four most generalist hummingbird species.

The NODF estimate also increased for the mite - plant network after including interactions with *Musa velutina* (NODF_original_= 29.9, NODF_novel_ = 30.8). This is the result of *M. velutina* interacting with the most generalist flower mite species, *Proctolaelaps* sp1.

We did not detect significant changes in relative nestedness between the original and novel hummingbird-plant networks after including the interactions with *M. velutina* (Permutation test, 1000 permutations, P = 0.58). There is no difference in network structure between the original and novel mite-plant networks (Permutation test, 1000 permutations, P = 0.81).

### Flight connections between native plants and *Musa velutina*

We identified a core group of five plant species connected to each other by four to five hummingbird species (Figure 2, group A). *Musa velutina* is included in the group of highly interconnected species. This core group is also connected with most plants in the community (Figure 2). A second group of plants share one to four hummingbird species amongst them (Figure 2, group B). In the third group, which includes most plant species (*i.e.* 41%), plants are connected by a single hummingbird species within the group (Figure 2, group C). A fourth group includes nine plant species missing direct interconnecting flights to more than half of the plant species (Figure 2, group D). Only one plant species, *Calathea crotalifera,* has no direct flight connections to *M. velutina* (Figure 2, group D)*. Erythrina cochleata* is the only plant species with not known flight connections with other host plants (Figure 2).

**Figure 2.**
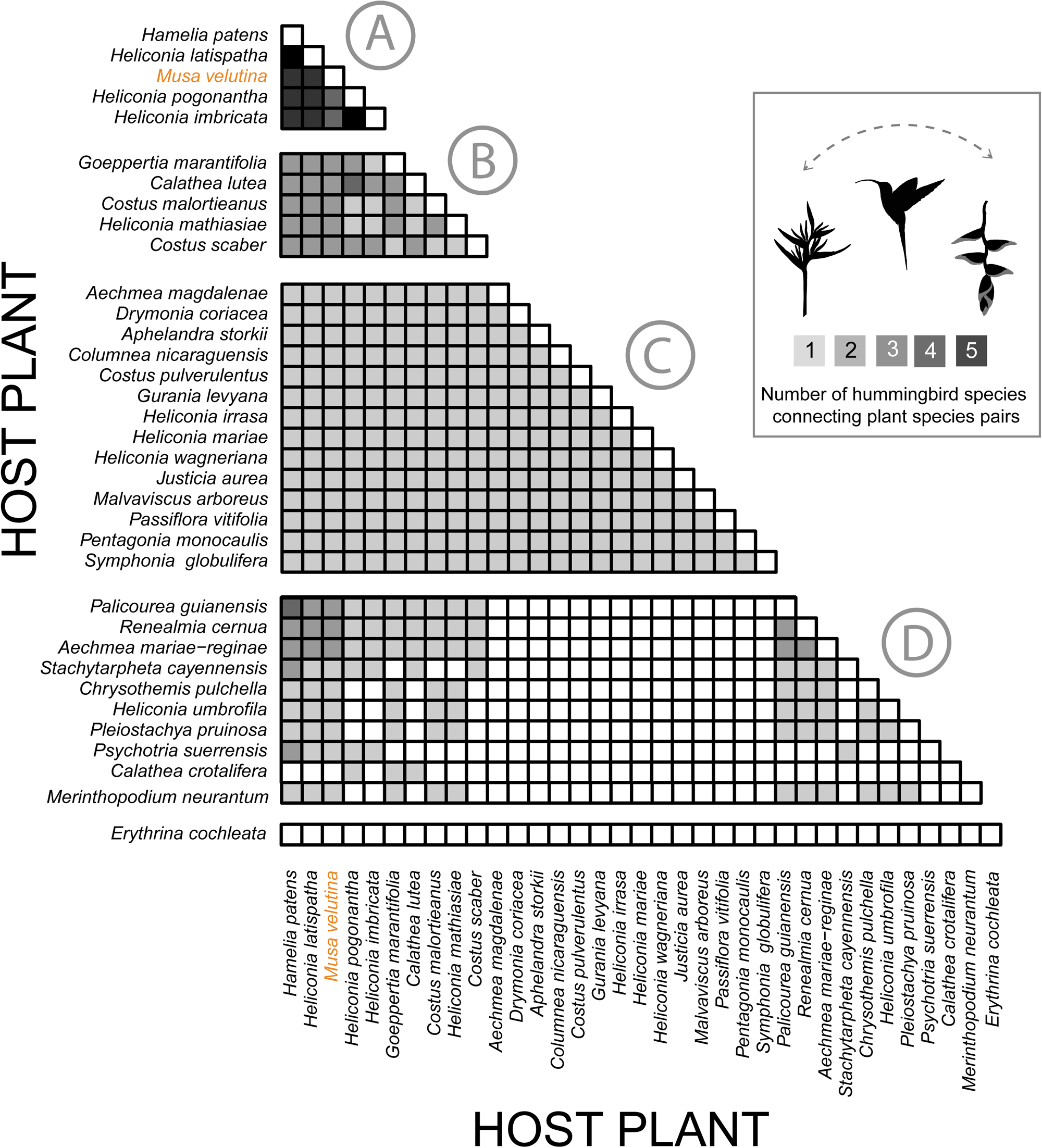
Flight connections between hummingbird-pollinated plants at La Selva. Each cell represents the number of hummingbird species connecting each pair of species. Groups include plant species with similar connectivity within the group. *Musa velutina* is highlighted in orange.

All flower mite species have the potential to take direct flights from at least one of their original hosts to *M. velutina* (Figure 3). From the 19 flower mite species recorded at La Selva, only *Proctolaelaps* sp1 disembarks in flowers of *M. velutina* (Figure 3). *Proctolaelaps* sp1 is the most generalist mite, and therefore the mite species with the highest number of direct flights to *Musa velutina* (Figure 3).

**Figure 3.**
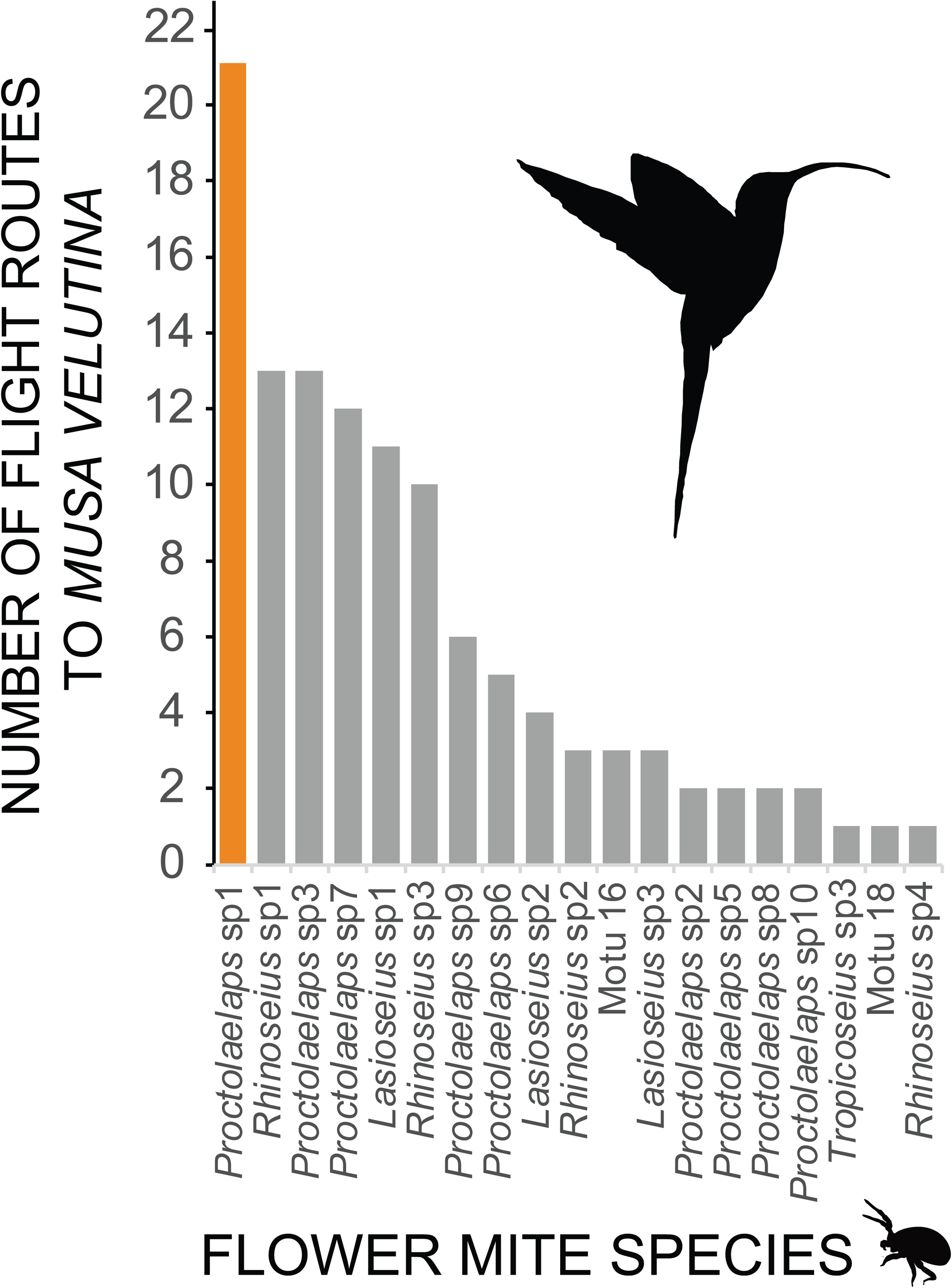
Number of flight connections, estimated as the sum of hummingbird species visiting each original host and *Musa velutina.* Mite species colonizing *Musa velutina* highlighted in orange.

### Hummingbird visits and flower mite colonization of *Musa velutina*

There is no difference in the number of hummingbird visits between female and male flowers (F_sex_ = 0.007, DF_sex_ = 25, P_sex_ = 0.09, F_plant_ = 1.22, DF_plant_ = 24, P_plant_ = 0.28 , Figure 4A). However, we detected a difference in the number of mites disembarking and embarking during hummingbird visits. In both female and male flowers, less than a half of the flower mites arriving to *M. velutina* eventually migrate to other inflorescences (F_sex_ = 0.0248, DF_sex_ = 40, P_sex_ = 0.87, F_transport_ = 5.49, DF_transportt_ = 24, P_transport_ = 0.02, Figure 4B). Male flowers harbor more mites than female flowers (F = 13.8, DF = 1, P = 0.0003). Male flowers have in average 15 times more mites than a female flower (Figure 4C).

**Figure 4.**
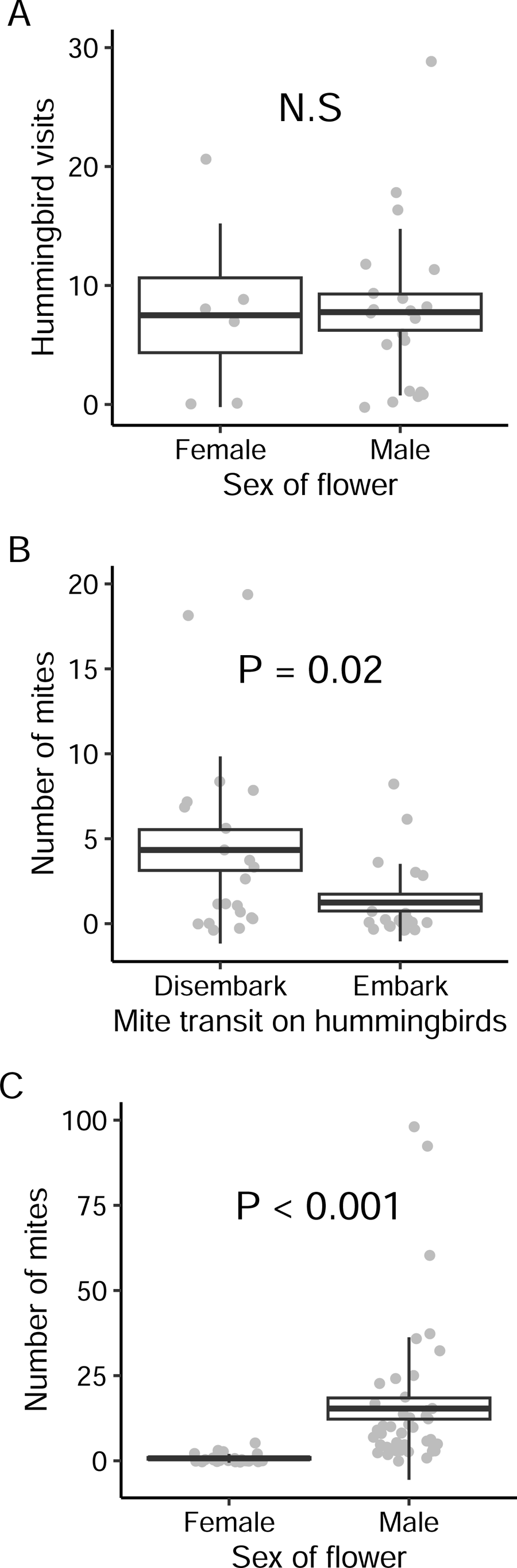
Hummingbird visits and mite colonization of female and male flowers of *Musa velutina* (Mean, SE and SD). A. Number of hummingbird visits to female or male flower hands. (N _flowers ♀_ = 36, N _flowers ♂_= 140). B. Number of mites disembarking or embarking from hummingbird beaks during each flower visit (N _hummingbird visits ♀_ = 66, N _hummingbird visits ♂_= 266). C. Number of flower mites inside female and male flowers (N _flowers ♀_ = 29, N _flowers ♂_ = 45).

## DISCUSSION

Our observations of generalist hummingbird species visiting *Musa velutina* represent another example of how hummingbirds usually incorporate novel hosts in their diets. In natural and urban settings, generalist hummingbirds tend to visit exotic plants, increasing the overall nestedness in the interaction network (Sánchez & Lara, 2024). Our study supports the notion that the ubiquitous nested structure in plant-pollinator networks is the result of generalist pollinators exploiting resources not visited by more specialized pollinators.

One common feature in plant-pollinator networks is the presence of keystone pollinator species, *i.e.*, species that serve as the only pollinators of many specialized plant species (Traveset, Tur, & Eguíluz, 2017). At La Selva, 40% of all hummingbird-pollinated plant species are only visited by *P. longirostris*. If this pollinator became locally extinct, these plants may lose their only potential pollinator. It is predicted that after pollinator extinctions, the survival of plant species specialized to a single pollinator will depend on their ability to self-pollinate or propagate vegetatively (Johnson & Steiner, 2000).

For the flower mite community, we found that 42% mite species specialize on a single host plant. However, these specialized species were recorded in six different host plants. This specialization in different plant species may reduce the risk of secondary extinction in flower mite communities (Maia, Marquitti, Vaughan, Memmott, & Raimundo, 2021; Traveset et al., 2017).

We recorded three plant species that are not hosts of flower mites. *Merinthopodium neurantum* is bat pollinated, and opens its greenish pendulous flowers at dusk (Bechler, Steiner, & Tschapka, 2024). The hummingbird *P. striigularis* visits *M. neurantum* flowers once, just before sunset. *Palicourea guianensis* and *Stachytarpetha cayennensis* open their flowers in the morning. Both plants are visited by hummingbirds during the day. More research is needed to determine which traits deter flower mites from colonizing these three plant species.

In this study, we were interested in determining if hummingbirds flight connections from native plants to *Musa velutina* facilitate or constraint the potential colonization of flower mites. Although all flower mite species have the potential to colonize *M. velutina*, only one of the nineteen flower mite species was recorded in the novel host plant. Hummingbirds at La Selva may carry multiple mite species in their beaks (Bizzarri, 2020). The two hummingbird species most frequently visiting *M. velutina, i.e.*, *P. longirostris*, and *A. tzacatl* may carry on their beaks as many as eight mite species, one of them is *Proctolaelaps* sp1 (Bizzarri, 2020). This shows that flower mites can select the plant species in which they disembark.

Previous studies in the laboratory reported that flower mites select their host plants using flower scents (Heyneman, Colwell, Naeem, Dobkin, & Hallet, 1991). These experiments might not be biologically relevant, as they report a very slow response of mites to floral scents. Mites selected their host after minutes or even hours of exposure to the scent cue (Heyneman et al., 1991). In a recent study we discovered that flower mites are almost instantaneously attracted to electric fields generated by hummingbirds (García-Robledo, Dierick, & Manser, Submitted). Because flowers are negatively charged, it is possible that flower mites can identify host plants by maybe combining electrical and chemical cues (García-Robledo et al., Submitted).

*Musa velutina* produces nectar with sugar concentrations within the range of those reported for native species (McDade & Weeks, 2004). Floral resources are scarce during the dry season (Stiles, 1975, García-Robledo, obs. pers.). Because *M. velutina* flowers all year, this exotic species might become a key resource for hummingbirds, and for at least one species of flower mite.

Although female flowers of *M. velutina* produce less nectar than male flowers, we did not observe any difference in the number of visits by hummingbirds. The number of mites disembarking in female and male flowers is also similar. This suggests that flower mites have similar probabilities of arrival to both female and male flowers. Colonization rates of *M. velutina* inflorescences are similar at both female and male phases. However, more mites were observed disembarking than embarking on hummingbird beaks. This suggests that only a fraction of the population of flower mites will eventually migrate to colonize novel hosts.

We recorded more flower mites in male than in female flowers. As shown in our previous results, the high abundance of flower mites in male flowers is not the result of intrinsic preferences of hummingbirds or flower mites (*e.g*., more hummingbirds visits or more mites disembarking in male flowers). The observed higher abundance of flower mites in male flowers is more likely the result of an accumulation of mites disembarking during hummingbird visits, and population growth during the time that inflorescences are producing flowers.

In conclusion, an exotic plant provides key services to pollinators and nectar parasites. *Musa velutina* seems to be fully integrated in the plant-pollinator network at La Selva, with the most generalist pollinators connecting this exotic host with most native plant species. This study provides additional evidence showing that hummingbird flower mites are not extremely specialized, as suggested by previous studies (Colwell, 1986). At least one flower mite species has the potential to colonize novel host plants through ecological fitting.

## DATA AVAILABILITY STATEMENT

Datasets available in Dryad, DOI: 10.5061/dryad.9kd51c5t1. Link for reviewers: http://datadryad.org/stash/share/4FlKicu2rAn2wTLoKg2phmVf2gCF3JCNebwhBI7ac3E.

## ACKNOWLEDGEMENTS

The authors thank the staff of La Selva Biological Station - Organization for Tropical Studies. We want to thank J.G. Huertas and J. Castro, for assistance in the field and laboratory. We thank D. Askew and L. Bizzarri for performing part of the molecular work. We are grateful to R. Bassini-Silva for identifying mite voucher specimens associated with CO1 sequences. Funding was provided by The National Science Foundation (Dimensions of Biodiversity 1737778). We thank the Comisión Nacional para la Biodiversidad (CONAGEBIO), for providing all required permits to perform this research (R-010-2024-OT). Comments by E.K. Kuprewicz and anonymous reviewers improved this manuscript.

## Notes

### Competing Interest Statement

The authors have declared no competing interest.

