## Supplement S1 for "Integration of an invasive plant in hummingbird and flower mite networks is driven by ecological fitting and generalization"

**Supplement S1.** Hummingbird-plant interactions at La Selva Biological Station, Costa Rica.

| **Family** | **Genus** | **Species** | **Hummingbird species** | **Reference** |
| --- | --- | --- | --- | --- |
| ACANTHACEAE | *Aphelandra* | *storkii* | *Phaethornis longirostris* | (McDade and Weeks 2004, Stiles and Wolf 1978) |
| ACANTHACEAE | *Justicia* | *aurea* | *Phaethornis longirostris* | (McDade and Weeks 2004, Stiles and Wolf 1978) |
| BROMELIACEAE | *Aechmea* | *magdalenae* | *Phaethornis longirostris* | (Stiles and Wolf 1978) |
| BROMELIACEAE | *Aechmea* | *mariae-reginae* | *Phaethornis striigularis* | Garcia-Robledo, video 14 Mar 2024 |
| CLUSIACEAE | *Symphonia* | *globulifera* | *Phaethornis longirostris* | (Sanfiorenzo et al. 2018) |
| CLUSIACEAE | *Symphonia* | *globulifera* | *Thalurania colombica* | J. Alvarado, obs. pers. |
| COSTACEAE | *Costus* | *malortieanus* | *Phaethornis longirostris* | (Kay & Schemske 2003, Stiles and Wolf 1978, Garcia-Robledo: Video) |
| COSTACEAE | *Costus* | *malortieanus* | *Phaethornis striigularis* | (Bizzarri 2020, Garcia-Robledo: Videos) |
| COSTACEAE | *Costus* | *pulverulentus* | *Phaethornis longirostris* | (Kay & Schemske 2003) |
| COSTACEAE | *Costus* | *scaber* | *Amazilia tzacatl* | (Kay & Schemske 2003) |
| COSTACEAE | *Costus* | *scaber* | *Phaethornis longirostris* | (Kay & Schemske 2003) |
| CUCURBITACEAE | *Gurania* | *levyana* | *Phaethornis longirostris* | (Stiles and Wolf 1978) |
| FABACEAE | *Erythrina cochleata* | *cochleata* | *Heliomaster longirostris* | (Neill 1987) |
| GESNERIACEAE | *Chrysothemis* | *pulchella* | *Paethornis striigularis* | Garcia-Robledo, obs. pers. 4 Mar 2024 |
| GESNERIACEAE | *Columnea* | *nicaraguensis* | *Phaethornis longirostris* | (Stiles and Wolf 1978) |
| GESNERIACEAE | *Drymonia* | *coriacea* | *Phaethornis longirostris* | (Stiles and Wolf 1978) |
| HELICONIACEAE | *Heliconia* | *imbricata* | *Amazilia tzacatl* | (Stiles 1975, Bizzarri 2020) |
| HELICONIACEAE | *Heliconia* | *imbricata* | *Chalybura urochrysia* | (Weathers and Styles 1989, Bizzarri 2020) |
| HELICONIACEAE | *Heliconia* | *imbricata* | *Glaucis aeneus* | (Bizzarri 2020, Garcia-Robledo obs. pers. 2024) |
| HELICONIACEAE | *Heliconia* | *imbricata* | *Phaethornis longirostris* | (Stiles 1975, Stiles and Wolf 1978, Bizzarri 2020) |
| HELICONIACEAE | *Heliconia* | *imbricata* | *Thalurania colombica* | (Linhart 1973, Stiles 1975, Bizzarri 2020) |
| HELICONIACEAE | *Heliconia* | *irrasa* | *Phaethornis longirostris* | (Stiles 1979, Bizzarri 2020) |
| HELICONIACEAE | *Heliconia* | *latispatha* | *Amazilia tzacatl* | (Stiles 1975, Bizzarri 2020) |
| HELICONIACEAE | *Heliconia* | *latispatha* | *Chalybura urochrysia* | (Stiles 1975) |
| HELICONIACEAE | *Heliconia* | *latispatha* | *Eutoxeres aquila* | J. Alvarado, obs. pers. |
| **Family** | **Genus** | **Species** | **Hummingbird species** | **Reference** |
| HELICONIACEAE | *Heliconia* | *latispatha* | *Phaethornis longirostris* | (Stiles and Wolf 1978) |
| HELICONIACEAE | *Heliconia* | *latispatha* | *Polyerata amabilis* | J. Alvarado, obs. pers. |
| HELICONIACEAE | *Heliconia* | *latispatha* | *Thalurania colombica* | (Stiles 1975, J. Alvarado obs. pers.) |
| HELICONIACEAE | *Heliconia* | *mariae* | *Phaethornis longirostris* | (Stiles and Wolf 1978) |
| HELICONIACEAE | *Heliconia* | *mathiasiae* | *Phaethornis longirostris* | (McDade and Weeks 2004, Bizzarri 2020) |
| HELICONIACEAE | *Heliconia* | *mathiasiae* | *Phaethornis striigularis* | (Bizzarri 2020, Garcia-Robledo obs. pers. 2023) |
| HELICONIACEAE | *Heliconia* | *pogonantha* | *Amazilia tzacatl* | (Bizzarri 2020, Garcia-Robledo, Video) |
| HELICONIACEAE | *Heliconia* | *pogonantha* | *Chalybura urochrysia* | (Stiles 1975, 1978, 1980) |
| HELICONIACEAE | *Heliconia* | *pogonantha* | *Eutoxeres aquila* | (Gill 1987, Stiles 1975) |
| HELICONIACEAE | *Heliconia* | *pogonantha* | *Glaucis aeneus* | (Bizzarri 2020) |
| HELICONIACEAE | *Heliconia* | *pogonantha* | *Phaethornis longirostris* | (Gill 1987, Stiles 1975, 1978, 1980, Stiles and Wolf 1978, Garrison 1995, Bizzarri 2020, Garcia-Robledo: Video) |
| HELICONIACEAE | *Heliconia* | *pogonantha* | *Thalurania colombica* | (Stiles and Wolf 1978, Bizzarri 2020) |
| HELICONIACEAE | *Heliconia* | *pogonantha* | *Threnetes ruckeri* | (Bizzarri 2020) |
| HELICONIACEAE | *Heliconia* | *sarapiquensis* | *Phaethornis longirostris* | (Bizzarri 2020)  removed from this study. Misidentification, red morph of *H. latispatha* |
| HELICONIACEAE | *Heliconia* | *sarapiquensis* | *Phaethornis striigularis* | (Bizzarri 2020)  removed from this study. Misidentification, red morph of *H. latispatha* |
| HELICONIACEAE | *Heliconia* | *umbrofila* | *Phaethornis striigularis* | (Stiles 1979) |
| HELICONIACEAE | *Heliconia* | *wagneriana* | *Phaethornis longirostris* | (Stiles and Wolf 1978, Garcia-Robledo: Video) |
| MALVACEAE | *Malvaviscus* | *arboreus* | *Phaethornis longirostris* | (Stiles and Wolf 1978) |
| MARANTACEAE | *Calathea* | *crotalifera* | *Threnetes ruckeri* | (Bizzarri 2020) |
| MARANTACEAE | *Calathea* | *lutea* | *Phaethornis longirostris* | (Stiles and Wolf 1978) |
| MARANTACEAE | *Calathea* | *lutea* | *Threnetes ruckeri* | (Stiles 1980) |
| MARANTACEAE | *Calathea* | *lutea* | *Amazilia tzacatl* | Garcia-Robledo, obs. pers. Mar 2024 |
| MARANTACEAE | *Calathea* | *marantifolia* | *Heliothryx barroti* | (Bizzarri 2020) |
| MARANTACEAE | *Calathea* | *marantifolia* | *Phaethornis longirostris* | (Bizzarri 2020) |
| MARANTACEAE | *Calathea* | *marantifolia* | *Phaethornis striigularis* | Garcia-Robledo pers. Obs. Nov 2023 |
| MARANTACEAE | *Calathea* | *marantifolia* | *Threnetes ruckeri* | (Stiles 1980, Bizzarri 2020, J. Alvarado, obs. pers.) |
| **Family** | **Genus** | **Species** | **Hummingbird species** | **Reference** |
| MARANTACEAE | *Pleiostachya* | *pruinosa* | *Phaethornis striigularis* | Kuprewicz, E.K. Video |
| PASSIFLORACEAE | *Passiflora* | *vitifolia* | *Phaethornis longirostris* | (Stiles and Wolf 1978, Gill et al. 1980) |
| RUBIACEAE | *Hamelia* | *patens* | *Amazilia tzacatl* | (Colwell 1995, Dearborn 1998, Lasso et al. 2003, Bizzarri 2020) |
| RUBIACEAE | *Hamelia* | *patens* | *Chalybura urochrysia* | (Lasso et al. 2003, Bizzarri 2020) |
| RUBIACEAE | *Hamelia* | *patens* | *Florisuga mellivora* | (Lasso et al. 2003) |
| RUBIACEAE | *Hamelia* | *patens* | *Heliomaster longirostris* | J. Alvarado, obs. pers. |
| RUBIACEAE | *Hamelia* | *patens* | *Klais guimeti* | (Dearborn 1998, Lasso et al. 2003, Bizzarri 2020) |
| RUBIACEAE | *Hamelia* | *patens* | *Lampornis alolaema* | (Lasso et al. 2003) |
| RUBIACEAE | *Hamelia* | *patens* | *Phaethornis longirostris* | (Bizzarri 2020, Garcia-Robledo obs. Pers. 2023) |
| RUBIACEAE | *Hamelia* | *patens* | *Phaethornis striigularis* | (Bizzarri 2020, Garcia-Robledo obs. Pers. 2023) |
| RUBIACEAE | *Hamelia* | *patens* | *Polyerata amabilis* | (Dearborn 1998, Lasso et al. 2003, Bizzarri 2020) |
| RUBIACEAE | *Hamelia* | *patens* | *Thalurania colombica* | (Colwell 1995, Dearborn 1998, Lasso et al. 2003, Bizzarri 2020, J. Alvarado pers. obs.) |
| RUBIACEAE | *Pentagonia* | *monocaulis* | *Phaethornis longirostris* | (Stiles and Wolf 1978) |
| RUBIACEAE | *Psychotria* | *suerrensis* | *Klais guimeti* | (Stone, 1994) |
| RUBIACEAE | *Psychotria* | *suerrensis* | *Thalurania colombica* | (Stone, 1994) |
| ZINGIBERACEAE | *Renealmia* | *cernua* | *Amazilia tzacatl* | (Bizzarri 2020, Carlos Garcia-Robledo obs. pers.) |
| VERBENACEAE | *Stachytarpheta* | *cayennensis* | *Amazilia tzacatl* | (Primack and Howe 1975) |
| VERBENACEAE | *Stachytarpheta* | *cayennensis* | *Klais guitmeti* | (Primack and Howe 1975) |
| ZINGIBERACEAE | *Renealmia* | *cernua* | *Phaethornis striigularis* | (McDade and Weeks 2004, Maglianesi et al. 2014, Bizzarri 2020) |

**Supplement S1**
